# Cerebrospinal fluid amyloid beta and glial fibrillary acidic protein concentrations in Huntington’s disease

**DOI:** 10.1101/2021.09.22.461351

**Authors:** Sara Korpela, Jimmy Sundblom, Henrik Zetterberg, Radu Constantinescu, Per Svenningsson, Martin Paucar, Valter Niemelä

## Abstract

**Introduction:** Huntington’s disease (HD) is a genetic incurable lethal disease. Biomarkers are needed for objective assessment of disease progression. Evidence supports both complex protein aggregation and astrocyte activation in HD. This study assesses the 42 amino acid long amyloid beta (Aβ42) and glial fibrillary acidic protein (GFAP) as potential biomarkers in the cerebrospinal fluid (CSF) of HD mutation carriers.

**Methods:** CSF was obtained from manifest HD patients (ManHD), premanifest HD-gene-expansion carriers (PreHD) and gene-negative controls (controls). Disease Burden Score (DBS) and Total Functional Capacity (TFC) were calculated. Protein concentrations were measured by enzyme-linked immunosorbent assays (ELISA) and intergroup differences were analysed using Mann-Whitney U test. Spearman correlations were calculated to assess disease stage association. Age-adjustment was included in the statistical tests.

**Results:** The study enrolled 27 ManHD and 13 PreHD subjects. The number of controls differed in the analysis of Aβ42 and GFAP (n = 19, and 8 respectively). Aβ42 levels were higher in ManHD (mean 741 ng/l, SD 361) compared with PreHD (mean 468 ng/l, SD 184) (p = 0.025). Likewise GFAP concentration was higher in ManHD (mean 435 ng/l, SD 255) compared with both PreHD (mean 266 ng/l, SD 92.4)(p = 0.040) and controls (mean 208 ng/l, SD 83.7)(p = 0.011). GFAP correlated with DBS (r = 0.361, p = 0.028), TFC (r = *−* 0.463, p = 0.005), and 5-year risk of onset in PreHD (r = 0.694, p = 0.008). In contrast, there was no correlation between Aβ42 concentration and DBS, TFC or 5-year risk of onset.

**Conclusion:** CSF Aβ42 levels did not correlate with disease stage suggesting no Aβaggregation in HD. GFAP is a potential biomarker in HD with association to disease stage. Validation in larger HD cohorts and potential correlations with clinical phenotype would be of interest.

## Introduction

Huntington’s disease (HD) is an autosomal dominant neurodegenerative disorder caused by pathological expansions in the polyglutamine (CAG expansion) region in exone 1 in the huntingtin gene (HTT) on chromosome 4q [1]. CAG encodes for the aminoacid glutamine (Q) and the CAG expansion leads to the formation of a protein with an abnormal poly-Q tail. Mutated HTT (mHTT) interacts with other proteins, accumulates and causes dysfunction and eventually degeneration and death of the neurons [2]. These changes have shown to be most prominent in capsula interna and striatum [3]. Symptoms of HD are progressive and include movement disorders, psychiatric symptoms [4] and cognitive decline [5]. There is no disease-modifying drug (DMD) or curative treatment available for HD so current treatment options are purely symptomatic. Several promising gene-based clinical trials are under way. HTT lowering therapies hold the promise of a disease modifiying therapy for HD. The diagnosis of HD is based on the presence of symptoms, clinical signs and confirmed by genetic testing. In the era of emerging therapies there is a need for biofluid markers to assess the progression of HD and thereby evaluate the utility of DMDs objectively. Some promising candidate biomarkers have recently been studied [6], of which neurofilament light protein (NfL), a broad biomarker for neurodegenerative diseases and neuroinflammation such as multple sclerosis (MS), seems to be the most promising with association to both disease onset and stage in HD [7–9], as well as mHTT [10]. Total tau (an axonal microtubule-stabilizing protein), another biomarker for neurodegeneration, has also been associated with the severity of symptoms, especially psychiatric symptoms but the data is less convincing compared with NfL [7, 9, 11–14]. Here we wanted to extend the CSF analysis of neurodegenerative markers with studies on Amyloid β(Aβ) protein and glial fibrillary acidic protein (GFAP) which have not been reported in HD before.

Dysregulation of amyloid β(Aβ) protein causes extracellular aggregation of the 42 amino acid form of Aβ(Aβ42) in brain tissue. Aβ42 is considered to be the most pathogenic form of Aβ [15]. Aβ42 aggregation leads to neuronal dysfunction mediated by excitotoxicity and neuroinflammation [16–18] and cereral hypoperfusion [19]. One form of Aβpathology is cerebral amyloid angiopathy (CAA), where the amyloid accumulates in the vascular walls [20]. Aβ42 aggregation is a hallmark of Alzheimer’s disease (AD) [21, 22], where CSF Aβ42 levels are reduced [23–26]. This decrease is likely due to increased amyloid deposition in the brain and decreased clearance into the CSF [27], and there is an inverse correlation with higher whole brain amyloid deposition load and low CSF Aβ42 [26, 28, 29]. Low Aβ42 levels may also be seen following axonal injury caused by e.g. traumatic brain injury [30], and in primary neurodegenerative disorders, such as dementia with Lewy bodies (DLB) [31] and Parkinson’s disease dementia [32]. CSF Aβ42 has been examined as a biomarker in depression where higher brain Aβburden in PiB-PET was found to be associated with increasing anxious-depressive symptoms over time in cognitively normal older individuals [33]. Studies on MS have found correlation between lower CSF Aβ42 levels and disease progression [34], including cognitive impairment [35], and higher gadolinium-enhancing lesion burden on magnetic resonance imaging [35].

GFAP (glial fibrillary acidic protein) is an intermediary fibrillary protein in the glial cells (astrocytes) and increased GFAP immunoreactivity is associated with gliosis and slowly progressing neuronal damage [36, 37]. It is used as a diagnostic marker in gliomas, especially astrocytomas [38, 39], is released in blood following ischaemic stroke [40], and is a marker for inflammation and disease stage in, e.g. MS [41–43]. Highly increased CSF GFAP concentration is seen in neuromyelitis optica spectrum disorders (NMOSD) due to rapid astrocytic damage [44, 45]. A post-mortem study performed on HD patients revealed astrocytosis in moderate, but not the earliest pathological stages of HD [46]. Similarly, trials on the HD mouse model R6/2 have shown increased GFAP levels in astrocytes of moderate and late disease stages. [47].

This study aims to investigate alterations of Aβ42 and GFAP levels as potential biomarkers for disease onset and progression in HD.

## Materials and methods

### Definition of participants and clinical assessment

The participants were recruited from the HD clinic at Uppsala University Hospital, from Karolinska University Hospital in Stockholm and Sahlgrenska University Hospital, Gothenburg, and were either premanifest gene-expansion carriers or manifest HD subjects. The gene-negative controls were recruited for the study amongst healthy individuals without signs of neurological disease. Premanifest gene-expansion carriers were defined as individuals with CAG expansion (over 35 repeats) of the HD-gene (HTT) and with a diagnostic confidence level (DCL) below 4. Manifest HD subjects were defined as individuals with a CAG-expansion (over 35 repeats) in the HD-gene and a DCL of 4. Disease burden score (DBS) was calculated for each HD patient using the formula (CAG repeat length-35.5) x age to estimate the cumulative HD pathology exposure. Total functional capacity (TFC) and corresponding disease stages (1-4) were assessed. The study was conducted in accordance with the declaration of Helsinki and was approved by the regional ethical review board in Uppsala, Sweden (DNR 2012/274). All participants signed an informed consent before study entry.

### CSF sample collection and handling

CSF was collected by lumbar puncture according to standardized protocol for procedure, materials and handling, but the time of the lumbar puncture and relation to meals varied. Polypropylene tubes and collecting vessels were used to avoid protein adsorption. The CSF was put on ice and centrifuged at 4 degrees Celsius and 1300 G for 10 minutes. The acellular proportion was pipetted off for storage at minus 70 degrees Celsius until the time of analysis.

### Biochemical analyses

CSF Aβ42 concentration was measured by INNOTEST enzyme-linked immunosorbent assay (ELISA) (Fujurebio, Ghent, Belgium). CSF GFAP concentration was measured using an in-house ELISA, as previously described in detail [48].

### Statistical analyses

Tests for normality of distribution included Shapiro-Wilk and inspection of histogram and the skewness statistics were applied. Age was approximately normally distributed, but the protein levels were not. Intergroup differences in protein levels were tested with non-parametric tests (Mann-Whitney U test). If there was deemed to be an association to age and/sex this variable was included as a covariant in a linear regression analysis model. We performed Spearman rank correlation with all gene-expansion carriers pooled into one group to assess the correlation between the protein levels and TFC as well as DBS. All statistical analyses were performed on cross-sectionally sampled data. The level of significance was defined by p-value less than 0.05. Statistical analyses were performed using SPSS Statistics Subscription Build 1.0.0.1461

## Results

The study enrolled 59 participants (mean age; standard deviation, range see Table 1 and Table 2).

**Table 1.** Characteristics of the study population; Aβ42. Age, CAG repeat length and Disease burden are presented as Mean (SD)

| Group | N | Age | Male:Female ratio | CAG | Disease burden |
| --- | --- | --- | --- | --- | --- |
| Total | 59 | 49.2 (16.1) | 29 : 30 | — | — |
| Controls | 19 | 48.5 (16.8) | 7 : 12 | — | — |
| Premanifest HD | 13 | 37.2 (11.5) | 7 : 6 | 43.6 (3.9) | 274.6 (75.8) |
| Manifest HD | 27 | 55.4 (14.4) | 15 : 12 | 43.5 (3.3) | 419.6 (187.7) |

**Table 2.** Characteristics of the study population; GFAP. Age, CAG repeat length and Disease Burden are presented as Mean (SD)

| Group | N | Age | Male:Female ratio | CAG | Disease burden |
| --- | --- | --- | --- | --- | --- |
| Total | 48 | 48.9 (16.2) | 28 : 20 | — | — |
| Controls | 8 | 42.9 (18.5) | 5 : 3 | — | — |
| Premanifest HD | 13 | 37.2 (11.5) | 7 : 6 | 43.6 (3.9) | 274.6 (75.8) |
| Manifest HD | 27 | 56.3 (13.6) | 16 : 11 | 43.6 (3.2) | 403.4 (202.9) |

CSF Aβ42 concentration was measured in 59 participants (mean age 49.2; SD 16.1, range 20 *−* 78 years). 13 of these (22.0%) were premanifest gene-expansion carriers, 27 (45.8%) manifest HD patients and 19 (32.2%) gene-negative controls.

CSF GFAP concentration was measured in 48 participants (mean age 48.9, SD 16.2, range 20 *−* 78 years). 13 of these (27.1%) were premanifest gene-expansion carriers, 27 (56.3%) manifest HD patients and 8 (16.7%) gene-negative controls. The characteristics of these subgroups are described in Table 1 and Table 2.

The protein levels were also compared to those of the laboratorys age-stratified reference rates. The group sizes differ because of insufficient amounts of CSF from several individuals, precluding quantitative analysis of both Aβ42 and GFAP.

### CSF Aβ42 and GFAP concentration differences between groups

There was no association between age and CSF Aβ42 concentration (r = − 0.322 p = 0.179) or CSF GFAP concentration (r = − 0.167, p = 0.693). Previous studies have not found a correlation between age and Aβ42 concentration in healthy individuals [49–51], but regarding GFAP a significant positive correlation between age and protein concentration has been noted in healthy individuals [48].

Regarding Aβ42, the premanifest and the manifest HD group differed significantly in age (p = 0.001), and regarding GFAP the manifest HD group was significantly older than both the premanifest HD group and control group (p = 0.0003 and p = 0.043 respectively), due to the natural course of the disease. To exclude potential confounding age was included as a covariate in the statistical analyses.

Figure 1 shows the concentrations of both proteins in the three groups. CSF Aβ42 concentration was significantly higher in the manifest HD group (mean 741 ng/l, SD 361) compared with the premanifest HD group (mean 468 ng/l, SD 183, (p = 0.025). The difference remained significant after adjustment for age (p = 0.03). There were no significant differences between the gene-negative controls (mean 535 ng/l, SD 238) and the manifest HD group, but a trend towards higher levels in the manifest HD group was noted (p = 0.076, age-adjusted p = 0.112). There were no significant differences between gene-negative controls and premanifest gene-expansion carriers (p = 0.454, age-adjusted p = 0.626).

**Fig 1.**
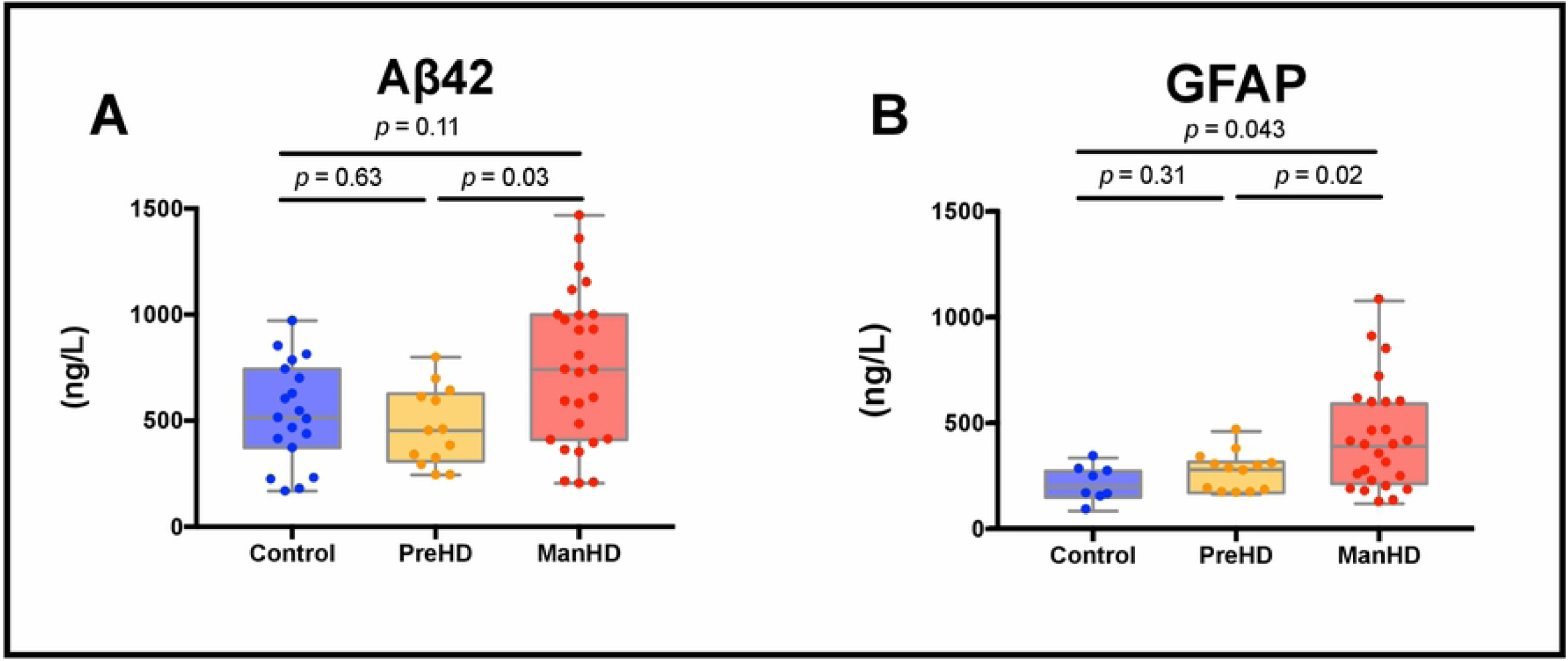
Concentrations of Aβ42 and GFAP by group. Protein concentrations plotted un-adjusted by group, p-values are adjusted for age. Boxes show first and third quartiles, the central bands show the median, and the whiskers show data within 1.5 IQR of the median. (A) Aβ42 levels were higher in the manifest group compared with the premanifest gene-expansion carriers. (B) GFAP levels were higher in the manifest group compared with both the premanifest gene-expansion carriers and healthy controls.

CSF GFAP concentration was significantly higher in the manifest HD group (mean 435 ng/l, SD 255) compared with both the premanifest HD group (mean 266 ng/l, SD 92.4)(p = 0.040) and the gene-negative controls (mean 208 ng/l, SD 83.7)(p = 0.011). These differences remained significant even after adjustment for age (p = 0.018, p = 0.043, respectively). CSF GFAP concentration did not differ significantly between the premanifest group and the gene-negative controls (p = 0.070, age-adjusted p = 0.313).

### Associations of CSF Aβ42 and GFAP concentrations with disease progression

As shown in Figure 2, we found no correlation between CSF Aβ42 concentration and DBS (r = 0.207, p = 0.220). Neither was there any correlation between Aβ42 concentration and TFC (r = *−* 0.270, p = 0.111). After adjustment for age the correlation between Aβ42 concentration and TFC became statistically significant (p = 0.024). Aβ42 concentration did not correlate with 5-year risk of onset among premanifest gene-expansion carriers (r = 0.218, p = 0.474).

**Fig 2.**
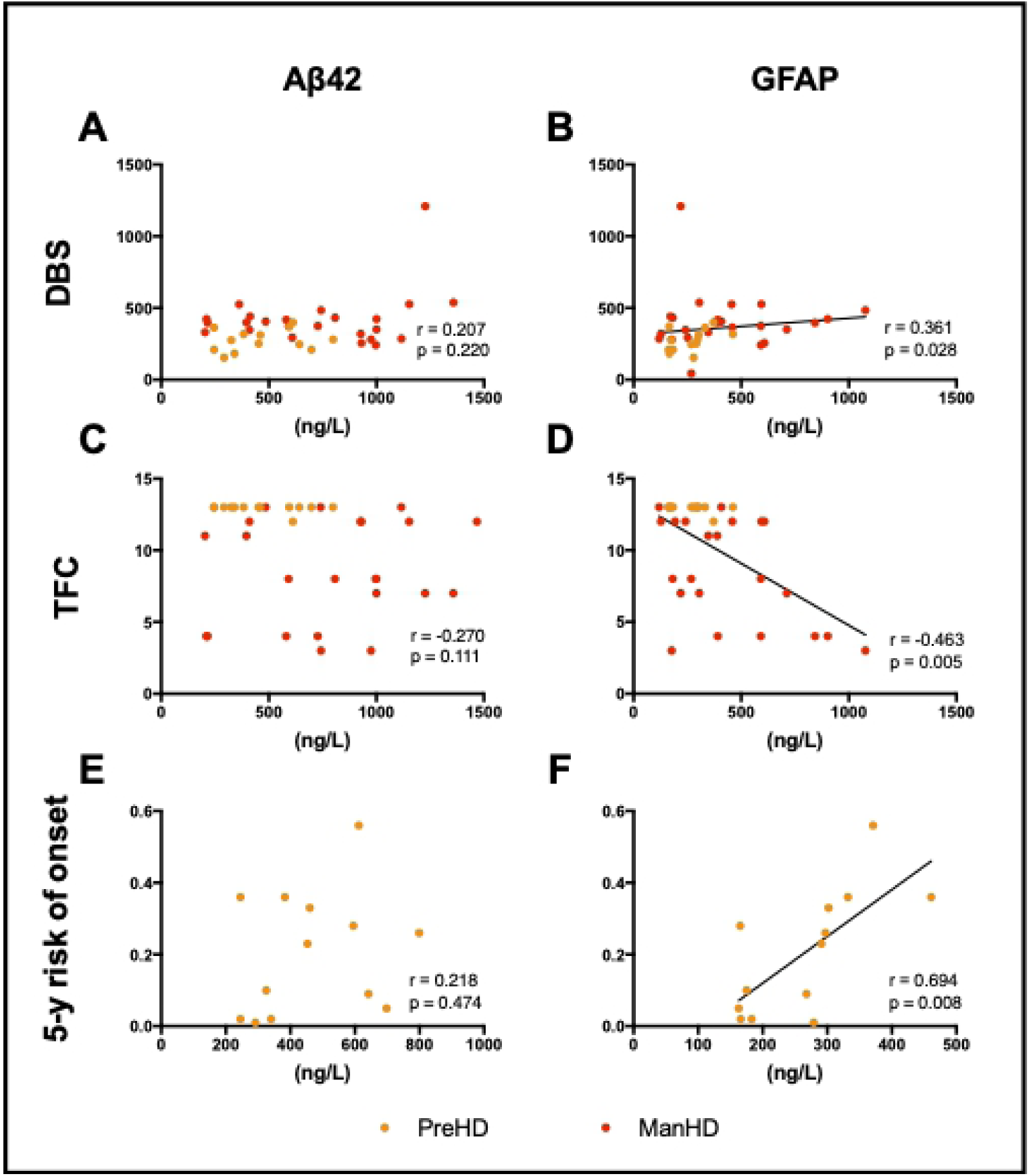
Correlations between protein levels and disease progression in Huntington’s disease. Aβ42 concentration does not correlate with (A) disease burden, (C) total functional capacity or (E) 5-year risk of onset, whereas GFAP concentration correlates with (B) Disease burden, (D) total functional capacity and (F) most strongly with 5-year risk of onset.

CSF GFAP concentration correlated positively with DBS and inversely to TFC with a significant correlation (r = 0.361, p = 0.028 and r = *−* 0.463, p = 0.005, respectively). Correlation between protein levels and TFC remained statistically significant even after adjustment for age (p = 0.001). DBS was not adjusted for age since this variable is already included in the composite score. GFAP levels correlated with 5-year risk of onset among premanifest gene-expansion carriers (r = 0.694, p = 0.008).

We did not find any correlation between CSF GFAP and Aβ42 concentrations (r =-0.097, p = 0.522).

## Discussion

In this exploratory CSF study we found significantly increased CSF Aβ42 concentration in the manifest HD group compared with the premanifest gene-expansion carriers. There was a non-significant tendency to lower levels of Aβ42 in the premanifest HD group compared with the healthy controls. However, Aβ42 concentration did not correlate with Disease Burden Score, Total Functional Capacity, or 5-year risk of onset.

A recent study demonstrated an association between lower levels of CSF amyloid precursor protein (APP) and worse clinical phenotype and lower cognitive performance in HD patients [52], the strongest relationship observed with composite UHDRS score. APP is a transmembrane protein with multiple physiological functions, including regulating brain iron homeostasis [53], and it is cleaved by beta- and gamma-secretase to form Aβ peptides [54, 55]. Our previous finding of decreased CSF transthyretin in HD patients [56] has also been reported in AD, where it possibly contributes to the failure of cerebral amyloid clearance [57]. This might allude to similarly decreased Aβ42 levels in HD, as noted in several other neurodegenerative disorders, most notably in AD [24–29]. On the contrary, we found the highest levels of Aβ42 in the manifest HD group. The mechanism underlying this finding is unknown. Nevertheless, the data corroborate that Aβ aggregation is not a feature of HD [58].

The present study found higher CSF GFAP concentration in the manifest HD group compared with both premanifest gene-expansion carriers and healthy controls. There was a non-significant trend towards higher CSF GFAP concentration in the premanifest group compared with controls. Further, GFAP concentration correlated positively with Disease Burden score and inversely with the Total Functional Capacity in the pooled gene-expansion carrier group. The strongest correlation was found between GFAP levels and 5-year risk of onset among the premanifest gene-expansion carriers. As GFAP is a marker of astrogliosis and degenerative process [36, 37], as well as a known marker for neuroinflammation [36, 37, 41–43], and/or astrocyte damage [44, 45], the interpretation of elevated GFAP levels is not so straightforward as to what kind of underlying pathological processes might be involved. The levels of GFAP also tend to rise with ageing in healthy individuals, most probably as a sign of astroglial filament formation in the CNS [48]. Still, our findings are in line with pathology studies of HD patients that have described astrocytosis in moderate, but not the earliest pathological grades of HD in humans [46], as well as findings in the R6/2 mouse, where GFAP was elevated in astrocytes of moderate and late disease stages as a sign of classical astrocyte activation [47]. Previous CSF studies in HD patients have shown increased CSF levels of YKL-40, another astrocytic activation marker, as a late feature of HD [9, 59], and a correlation to several markers of neurodegeneration [59]. YKL-40 is secreted by astrocytes and is increased in many inflammatory CNS disorders [60]. Evidence from CSF studies in HD mainly supports activation of the innate immune system of the CNS. This may reflect inflammation caused by neurodegeneration in the later stages of the disease [9]. However, an early increase of IL-6 and IL-8 levels in both CSF and plasma suggest an innate immune response both centrally and peripherally in HD [61]. There is also evidence of T-cell mediated inflammation ahead of disease onset [59]. The tendency to elevation of GFAP in premanifest gene-expansion carriers together with a strong correlation with risk of onset suggests astrocyte activation predating phenoconversion. Some of the premanifest gene-expansion carriers were far from predicted disease onset and this could be a reason that GFAP was not clearly elevated in this group.

To the best of our knowledge, this is the first CSF study performed on humans to assess the role of Aβ and GFAP as potential biomarkers in HD. However astrocyte activation in the form of astrocytosis and neuroinflammation has previously been linked to the pathophysiology of HD, especially in the late disease stages so this is in line with the finding of elevated GFAP levels amongst the manifest HD patients. Part of the elevation might be due to the normal aging process, as the manifest HD patients are usually older compared to the premanifest HD-gene-expansion carriers, but the difference remained significant even after taking this fact into account. The finding of the highest Aβ42 levels in the manifest HD group was unexpected considering that in other neurodegenerative diseases, Aβ tends to decrease in CSF along with the level of neurodegeneration in parallel with worse clinical outcome. Limitations in this study include the exploratory nature and a small sample size. Different ages between groups was of concern and the fact that the groups differed in size. The medications used by the patients and their potential effect on the protein levels was not assessed. Nonetheless, we believe that these findings may be of relevance regarding the involvement of astrocyte activation, neuroinflammation and gliosis in HD. GFAP may have a role in assessing the severity of HD, and could potentially also serve as a surrogate end-point in clinical trials. Before taking these findings into clinical practice there is a need for validation in a larger HD cohort and assessment of correlations with clinical phenotype would be of interest.

## Supporting Information

**S1 Table. Dataset including clinical characteristics of study participants and protein concentrations**. (PDF)

## Acknowledgments

We want to thank the participants of this study for donating their samples

## Author Contributions

**Planning:**

VN, JS

**Conceptualization:**

VN, JS, RC, PS, MP

**Data collection:**

VN, JS, RC, PS, MP, SK, HZ

**Formal analysis:**

HZ, SK

**Funding acquisition:**

JS, VN, HZ, PS

**Investigation:**

VN, HZ, SK

**Methodology:**

VN, SK, JS, PS

**Project administration:**

VN, RC, PS, JS, SK

**Software:**

SK, VN

**Visualization:**

VN, SK

**Writing - original draft:**

SK, VN, JS

**Writing - review & editing:** SK, VN, JS, HZ, RC, PS, MP

